# Discovery of NOvel CIP2A VAriant (NOCIVA) and its clinical relevance in myeloid leukemias

**DOI:** 10.1101/2020.08.24.264606

**Authors:** Eleonora Mäkelä, Karolina Pavic, Taru Varila, Urpu Salmenniemi, Eliisa Löyttyniemi, Srikar Nagelli, Veli-Matti Kähäri, Richard E Clark, Venkata Kumari Bachanaboyina, Claire Lucas, Maija Itälä-Remes, Jukka Westermarck

**Author notes:** Corresponding author: Jukka Westermarck.

## Abstract

Cancerous inhibitor of PP2A (CIP2A) is a prevalent human oncoprotein that inhibits tumor suppressor PP2A-B56a. However, *CIP2A* mRNA and protein variants remain uncharacterized. Here, we report discovery of a *CIP2A* splicing variant *NOCIVA* (NOvel CIp2a VAriant). *NOCIVA* contains *CIP2A* exons 1-13 fused to a continuous stretch of 349 nucleotide from *CIP2A* intron 13. Intriguingly, the first 39 nucleotides of the *NOCIVA* specific sequence are in coding frame with exon 13 of *CIP2A*, and codes for a 13 amino acid peptide tail unhomologous to any known human protein sequence. Therefore, NOCIVA translates to a unique human protein. NOCIVA retains the capacity to bind to B56a, but whereas CIP2A is predominantly a cytoplasmic protein, NOCIVA translocates to nucleus. Indicative of prevalent alternative splicing from *CIP2A* to *NOCIVA* in myeloid malignancies, acute myeloid leukemia (AML) and chronic myeloid leukemia (CML) patient samples overexpress *NOCIVA*, but not *CIP2A* mRNA. In AML, high *NOCIVA* mRNA expression is a marker for adverse overall survival. In CML, high *NOCIVA* expression associates with inferior event free survival among imatinib treated patients, but not among patients treated with dasatinib or nilotinib. Collectively, we describe discovery of a novel variant of oncoprotein CIP2A, and its clinical relevance in myeloid leukemias.

**Key Points:**

- Discovery and characterization of a first mRNA variant of one of the most prevalently deregulated human oncoproteins CIP2A
- Unlike CIP2A, NOCIVA mRNA is overexpressed in AML and CML patient samples and associates with poor clinical response in both myeloid cancers

## Introduction

Cancerous inhibitor of protein phosphatase 2A (CIP2A) functions as an oncoprotein by directly binding to tumor suppressor PP2A-B56α [1, 2]. CIP2A is overexpressed in a vast variety of human cancers, and high CIP2A expression has been shown to correlate with poor patient survival in a broad spectrum of human malignancies [3-5]. Furthermore, CIP2A is required for malignant cellular growth *in vitro* and for tumor formation *in vivo* in a number of cancers, and its overexpression promotes broadly cancer cell drug resistance [3, 5-10]. However, CIP2A deficiency does not compromise normal mouse development or growth [6, 11, 12]. Consequently, inhibition of CIP2A protein expression or activity could constitute a very efficient cancer therapy strategy with minor side effects. Prior to advancement of this potential cancer therapy target in drug development, and for the design of highly specific therapeutics, there should be a comprehensive understanding of CIP2A protein and/or mRNA variants. In regard to the *CIP2A (KIAA1524)* gene, there are no genetic homologs in the human genome, and apart from *CIP2A* variant database predictions that lack experimental validation, virtually no information exists about variant forms of CIP2A at either mRNA or protein level.

Alternative splicing (AS) is a physiological phenomenon that greatly diversifies the repertoire of the transcriptome. As up to 95% of multi-exonic genes are alternatively spliced [13], AS ensures higher protein diversity for better environmental fit. Perturbation of AS by spliceosome gene mutations, epigenetic modifications, or other causes lead to aberrant AS, and this has been shown to be a hallmark of cancer development [14]. Pan-cancer studies have revealed that tumors have an average of 20% more AS events than healthy samples [15, 16]. Current evidence suggests a pivotal role of AS abnormalities especially in leukemia pathogenesis, particularly in myelodysplastic syndrome and acute myeloid leukemia (AML) [17, 18].

Myeloid leukemias, including AML and chronic myeloid leukemia (CML), are heterogeneous clonal hematological malignancies that disrupts normal hematopoiesis. Whereas AML is the most common acute leukemia affecting adults, CML accounts for 15–25% of all adult leukemias [19, 20]. Interestingly, both AML and CML are one of the very few human malignancies in which *CIP2A* mRNA is not overexpressed, although presumably due to post-transcriptional stabilization, CIP2A is overexpressed at protein level and this correlates with more aggressive disease [21, 22]. Despite therapeutic progress, the outlook for AML remains unsatisfactory [23] and up to 50% of AML patients will relapse [23, 24]. On the contrary, CML treatment was revolutionized by the use of targeted tyrosine kinase inhibitors (TKIs), which have dramatically improved long-term survival [20, 25]. Particularly in AML, a prominent component of the disease is the recurrent mutations in spliceosome machinery and genome-wide aberrant splicing events [26]. As AS is an important part of normal hematopoiesis and necessary for cellular differentiation [27], the role of abnormal AS in leukemia progression and drug resistance has gained attention as several recent studies have highlighted recurrent splice factor mutations as important drivers of hematological malignancies [28-30].

Here we identified a novel CIP2A variant, NOCIVA (<u>NO</u>vel <u>CI</u>P2A <u>VA</u>riant), that is produced via AS. *NOCIVA* translates to a unique human protein that can heterodimerize with CIP2A and bind to PP2A-B56a subunit. Interestingly, NOCIVA is predominantly a nuclear protein. We also show that the expression of *NOCIVA* is elevated in cancer and that in myeloid cancers, such as AML and CML, high *NOCIVA* expression is a marker of a poor clinical outcome. Of particular clinical relevance, in CML high *NOCIVA* expression is associated with resistance to first generation TKI imantinib, but this effect is not seen with patients treated with second generation TKIs, dasatinib or nilotinib.

## Methods

See supplemental information for more detailed methods.

### 3’ RACE and 5’ RACE

For both 3’ and 5’ rapid amplification of cDNA ends, Invitrogen’s (Carlsbad, CA, USA) 3’RACE (catalog no. 183743-019) and 5’RACE (catalog no. 18374-058) kits were used according to the manufacturer’s protocols. Details in Supplemental Methods.

### Quantitative real-time PCR (RQ-PCR)

The standard curve analysis for amplification efficiency and the melting curve analysis for NOCIVA#1 and NOCIVA#2 RQ-PCRs are shown in Supplemental Figure 1. The primer and probe sequences used in this study for RQ-PCR analysis are listed in Supplemental Table 1. Details in Supplemental Methods.

**Figure 1.**
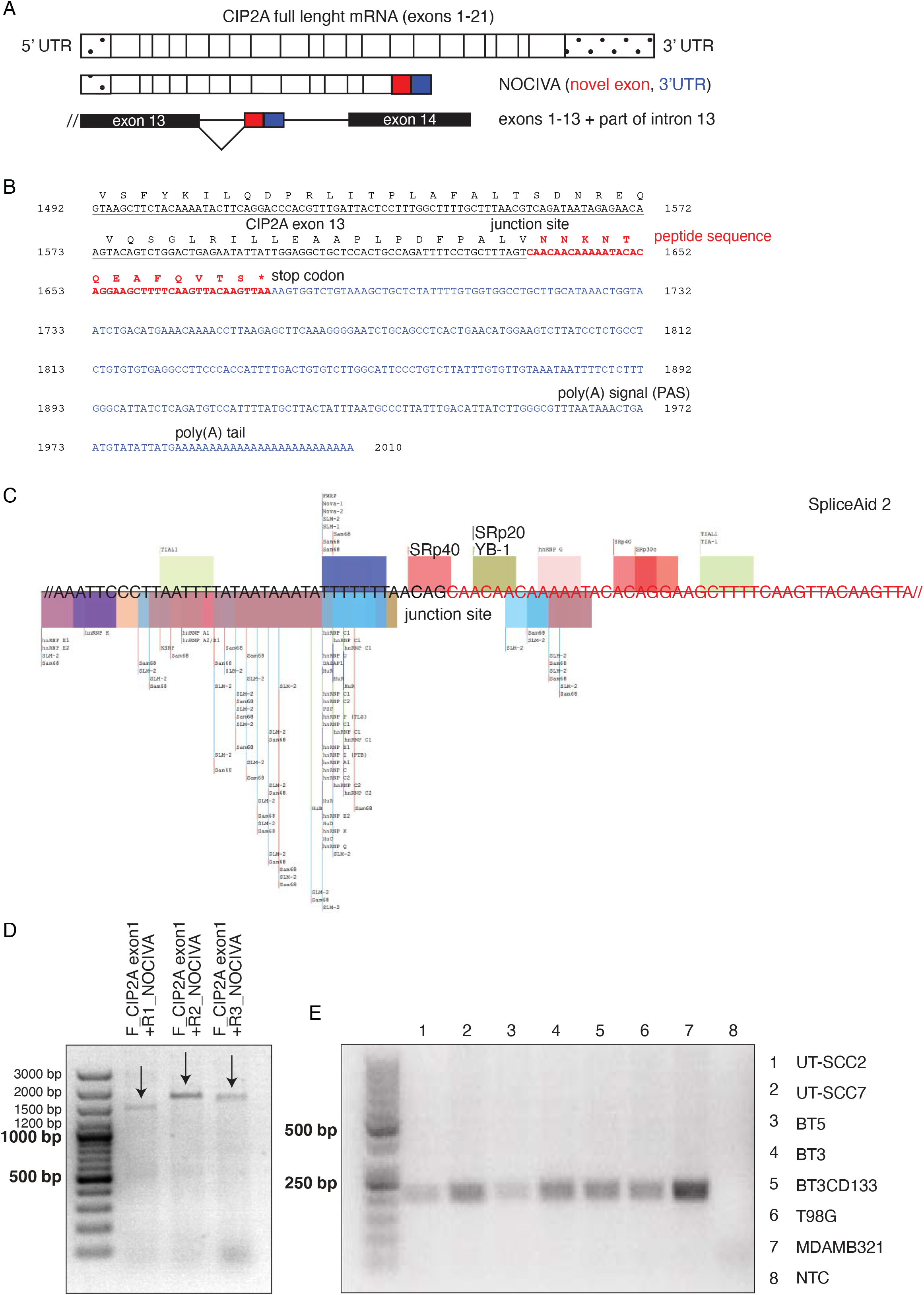
Characterization of Novel CIP2A Variant (NOCIVA) mRNA isoform. **A)** A schematic presentation of the *NOCIVA* mRNA isoform identified with RACE-PCR. *NOCIVA* mRNA contains an alternative exon from *CIP2A* intron number 13, and thus forms a unique and previously unknown coding sequence. Untranslated regions (5’UTR or 3’UTR) are marked with dots, the unique alternative exon in *NOCIVA* with red and *NOCIVA* specific 3’UTR with blue. Full length *CIP2A* stands for RefSeq NM_020890.2 sequence. **B)** *NOCIVA* mRNA’s 3’-end with the differing features from the original *CIP2A* mRNA sequence. The shared nucleotide sequence between *CIP2A* and *NOCIVA* mRNA is underlined. NOCIVA protein comprises 545 N-terminal CIP2A amino acids and 13 unique amino acids on the C-terminus (in bold and red). The stop codon is indicated by an asterisk. *NOCIVA* mRNA contains a 3’UTR (blue, 1675-2010) with a polyadenylation signal (PAS, AATAAA at 1962-1967). **C)** The *NOCIVA* splice junction with splice site predictions from SpliceAid 2 and SFmap. At the *NOCIVA* junction site there are binding motifs for YB-1, SRp20 (SRSF3) and SRp40 splicing factors. **D)** Confirmation PCR to validate the *NOCIVA* specific full-length mRNA sequence expression from the Hela cell line. The forward primer for all lanes was the same *CIP2A* exon1 targeting, the reverse primer was *NOCIVA* specific and either R1, R2 or R3. Arrows indicating the right product. Resulting bands were extracted and the presence of specific *NOCIVA* cDNA (mRNA) was confirmed by DNA sequencing. **E)** Confirmation PCR to validate the *NOCIVA* specific mRNA sequence expression from several cell lines with *CIP2A* exon13 targeting forward primer and the *NOCIVA* specific reverse primer. NTC stands for non-template control.

### Binding assay

Protein expression, purification and binding assays were performed as in [1]. Details in Supplemental Methods.

### NOCIVA antibodies

Two NOCIVA specific antibodies were generated by immunizing rabbits against NOCIVA specific peptide NNKNTQEAFQVTS by BioGenes GmbH (Berlin, Germany). Details in Supplemental Methods.

### Patient cohorts

#### AML study cohort

Detailed information for AML study cohort can be found in [31]. All 80 patients received regimens comprising anthracycline and high-dose cytarabine as induction therapy. Their median age was 50 years (Q_1_ = 38.8, Q_3_ = 58.0), median overall survival was 5.4 years (95% CI, 2.8 to 7.9) and median follow-up time was 5.4 years (range 6 days–16 years). The European LeukemiaNet (ELN) 2010 risk group classification [32] was used for risk stratification (Table S2).

#### CML study cohort1

This cohort comprised of 35 newly diagnosed chronic phase CML patients from the University of Liverpool CML biobank. One patient lacked follow up data. Twenty patients received imatinib as a first line therapy and 14 received a second generation TKI, either dasatinib or nilotinib. Their median age was 53.5 years (Q_1_ = 42.3, Q_3_ = 62.0), the median follow-up time was 32.5 months (range 9–75 months) and median event free survival was 30.9 months (95% CI, 24.1 to 39.4).

#### CML study cohort2

This cohort consisted of 159 newly diagnosed CML patients from the UK-wide SPIRIT2 clinical trial [33]. The samples were the first 141 biobanked samples plus 18 additional patients whose disease progressed. Eighty-one patients received imatinib and 78 dasatinib as their first line treatment. Their median age was 53 years (Q_1_ = 43, Q_3_ = 63) and median follow-up time was 60 months (range 1–60 months).

### Statistical analysis

Statistical analysis was performed using SAS software (version 9.3, SAS Institute Inc., Cary, NC, USA) or GraphPad Prism (version 8.3., GraphPad Software, San Diego, CA, USA). Normal distribution of the data was tested and if needed transformations were performed. All statistical tests were two-sided and declared significant at a p-value of less than 0.05. Details in Supplemental Methods.

### Data sharing statement

For original data, please contact.

## Results

### Identification of Novel CIP2A Variant (NOCIVA) mRNA isoform

To identify potential mRNA variants of *CIP2A* (gene alias *KIAA1524*), rapid amplification of cDNA ends PCR assays (3’RACE and 5’RACE) were employed in human cell line mRNA samples (PNT2, MDA-MB-231, HeLa). As expected from database searches (NCBI databases, Ensembl), one of the observed variants contained CIP2A exons 1-19, instead of 21 exons in the full length *CIP2A*. In addition, a novel CIP2A mRNA splice variant (named here as *NOCIVA*) with alternative exon inclusion was identified (Fig 1A). *NOCIVA* comprised of exons 1-13 of *CIP2A* fused C-terminally to a part of the intron between exons 13 and 14 (Fig 1A). This 349 nucleotide intronic region (Fig 1A, Fig S2B) is normally located inside the intron 13 of the *CIP2A* gene, more precisely ranging from 108561721 to 108562069 in Homo sapiens chromosome 3 (GRCh38.p13 reference, annotation release 109.20200228). As a clear evidence that NOCIVA constitutes a functional mRNA transcript, *NOCIVA* mRNA contains a stop codon followed by a 330 nucleotide 3’UTR with polyadenylation signal (PAS) AATAAA and poly(A) tail (Fig 1B and Fig S2A).

As an indication that *NOCIVA* mRNA is created by AS, the spliced intron region is flanked by GT and AG dinucleotides (Fig S2B yellow, GU-AG intron). Further, based on *in silico* analysis with Human Splicing Finder (version 3.1, [34]), the junction site between *CIP2A* and *NOCIVA* contains exonic splicing silencer (ESS) matrices, especially Fas ESS and PESS-octamers. ESSs work by inhibiting the splicing of pre-mRNA strands or promoting exon skipping. On the other hand, SpliceAid 2 [35] and SFmap (version 1.8, [36]) gave identical predictions for binding of YB-1 and SRp20 (SRSF3) splicing factors at the *NOCIVA* junction site (Fig 1C). Both of these splice factors have been shown to promote exon-inclusion during alternative splicing [37, 38]. Further, binding sites for many other splice factors, including Sam68, SLM2, SRp40 and multiple hnRNPs (including hnRNP K), were found in the near vicinity of the junction site (Fig 1C).

Validation PCR for full length *NOCIVA* mRNA expression was conducted in HeLa cell line with forward primer targeting *CIP2A* exon1 combined with various reverse primers targeting the *NOCIVA* specific 3’ end of the mRNA (Fig 1D and Fig S3A). Additionally, validation PCR for *NOCIVA* expression was conducted in multiple cancer cell lines with primers specific for mRNA coding for the unique C-terminal portion of *NOCIVA* (Fig 1E and Fig S3B). The correct size bands from the gels were subsequently sequenced to confirm that the PCR product represented the *NOCIVA* mRNA product.

Together, these results identify *NOCIVA* as a novel, alternatively spliced *CIP2A* variant that is expressed in multiple cancer cell lines.

### Characterization of NOCIVA protein

Interestingly, in *NOCIVA* mRNA, the 5’end of the *NOCIVA* specific intronic sequence is fused in coding frame with the preceding 3’end of the *CIP2A* mRNA sequence. After 40 nucleotides, corresponding to 13 amino acids (aa) (red text in Fig 1B) the C-terminal tail is followed by a classical stop codon TAA. Therefore, the potential NOCIVA protein consists of 545 aa that are shared with CIP2A, followed by the NOCIVA specific peptide sequence NNKNTQEAFQVTS (Fig 1B). The novel 13 aa peptide sequence in NOCIVA did not match with any known protein sequence in the human proteome based on Blast homology search [39] (Fig S4A, BLASTP 2.8.1+, Database: Non-redundant protein sequences (nr)). Next, we used the recombinant NOCIVA peptide to generate two affinity chromatography purified NOCIVA specific antibodies. The specificity of the antibodies was tested by using bacterially produced NOCIVA and CIP2A proteins. Anti-NOCIVA antibodies specifically recognized NOCIVA but neither full length CIP2A nor CIP2A protein fragments (Fig 2A, Fig S4B for NOCIVA ab #2 data). Importantly, the NOCIVA signal could be abolished by using blocking peptide (Fig 2A). To study spatial expression of endogenous NOCIVA, we performed immunofluorescence (IF)-staining in MDA-MB-231 breast cancer cells. Interestingly, whereas CIP2A resided predominantly in the cytoplasm as expected [3], endogenous NOCIVA positivity was clearly nuclear (Fig 2B and Fig S4C). Similar conclusion could be drawn from GFP fusion overexpression studies, in which NOCIVA-GFP colocalized with DAPI to nucleus (Fig 2C). With this approach also cytoplasmic NOCIVA-GFP was detected which was probably due to prominent colocalization of empty GFP to cytoplasm (Fig 2C). Hence, NOCIVA expresses a novel immunogenic peptide sequence, and constitutes a predominantly nuclear CIP2A variant protein.

**Figure 2.**
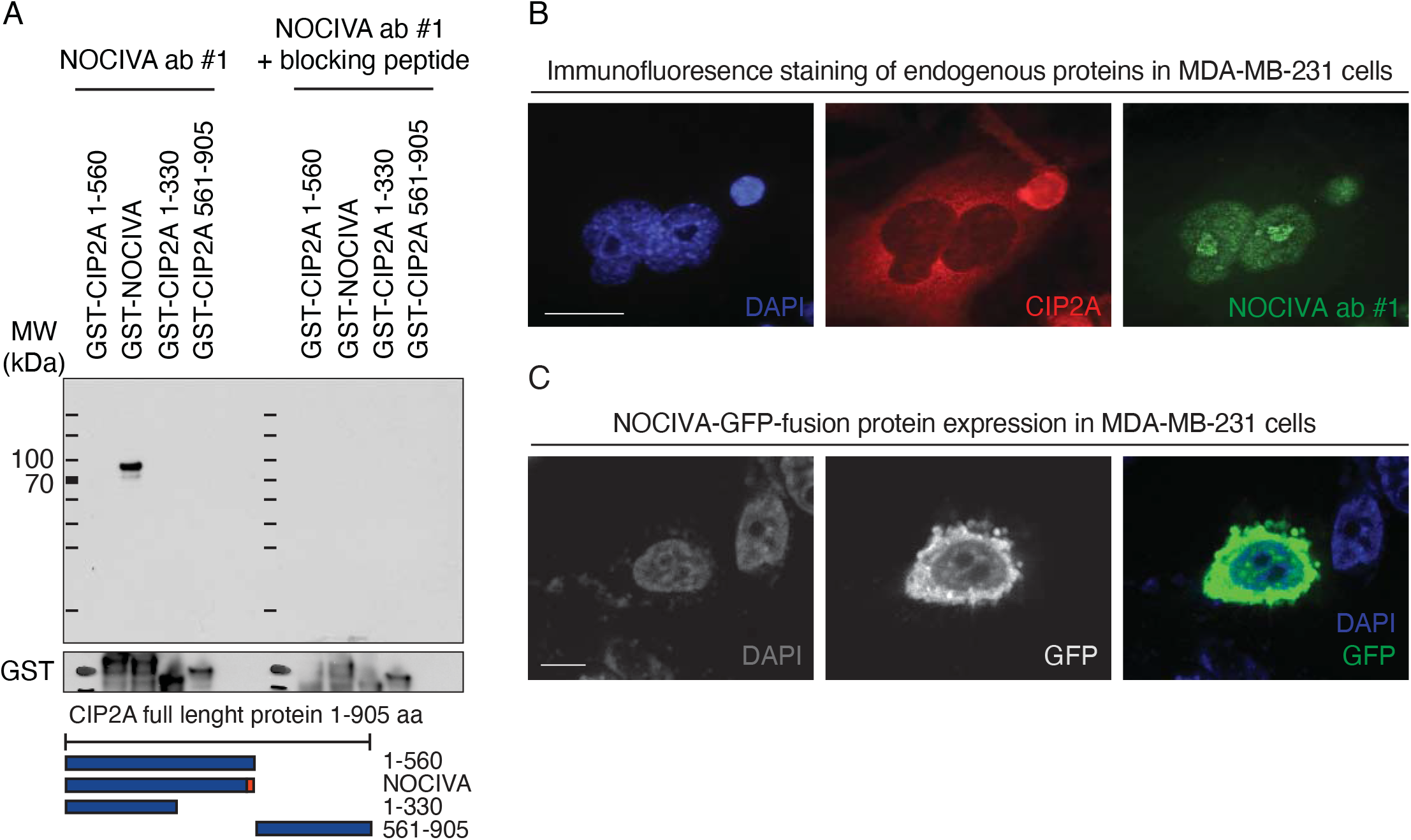
Characterization of NOCIVA protein in the cells. **A)** NOCIVA specific antibody detects correct size (appr. 90 kDa) recombinant GST-NOCIVA protein, but not recombinant CIP2A fragments. 1ug of each protein was loaded onto each gel. The signal is blocked with a NOCIVA-blocking peptide. Full length CIP2A comprises of 905 aa. **B)** Endogenous NOCIVA and CIP2A IF staining with NOCIVA specific antibody in MDA-MB-231 cells. NOCIVA green, CIP2A red, nucleus blue. Endogenous NOCIVA localizes mostly in the nucleus whereas full length CIP2A is mainly cytoplasmic. Scale bar 10 μm. **C)** NOCIVA-GFP overexpression in MDA-MB-231 cells. As seen with endogenous NOCIVA IF staining, GFP tagged NOCIVA also translocates to the nucleus. NOCIVA-GFP green, nucleus blue. Scale bar 10 μm.

To address NOCIVA protein functions, recombinant GST-NOCIVA (CIP2A 1-545+13 aa peptide) and GST-CIP2A 1-560 were compared (Fig S4D for Coomassie staining) in two functions critical for CIP2A-mediated PP2A modulation; protein homodimerization, and direct binding to the B56α subunit of PP2A [1]. Consistently with the location of the B56α binding regions in the N-terminal part of CIP2A 1-560 [1], which is identical between NOCIVA and CIP2A, both proteins co-immunoprecipitated B56α with equal efficiency *in vitro* (Fig 3A). Additionally, NOCIVA was competent to heterodimerize with CIP2A 1-560, albeit with lower affinity than that was seen with CIP2A 1-560 homodimers (Fig 3B). This can be explained as the CIP2A-NOCIVA fusion site partly overlaps with the aa region mediating CIP2A homodimerization [1](Fig 3C,D) and that when compared to CIP2A homodimers, in NOCIVA-CIP2A heterodimers some of the stabilizing interactions are lost (Fig 3E).

**Figure 3.**
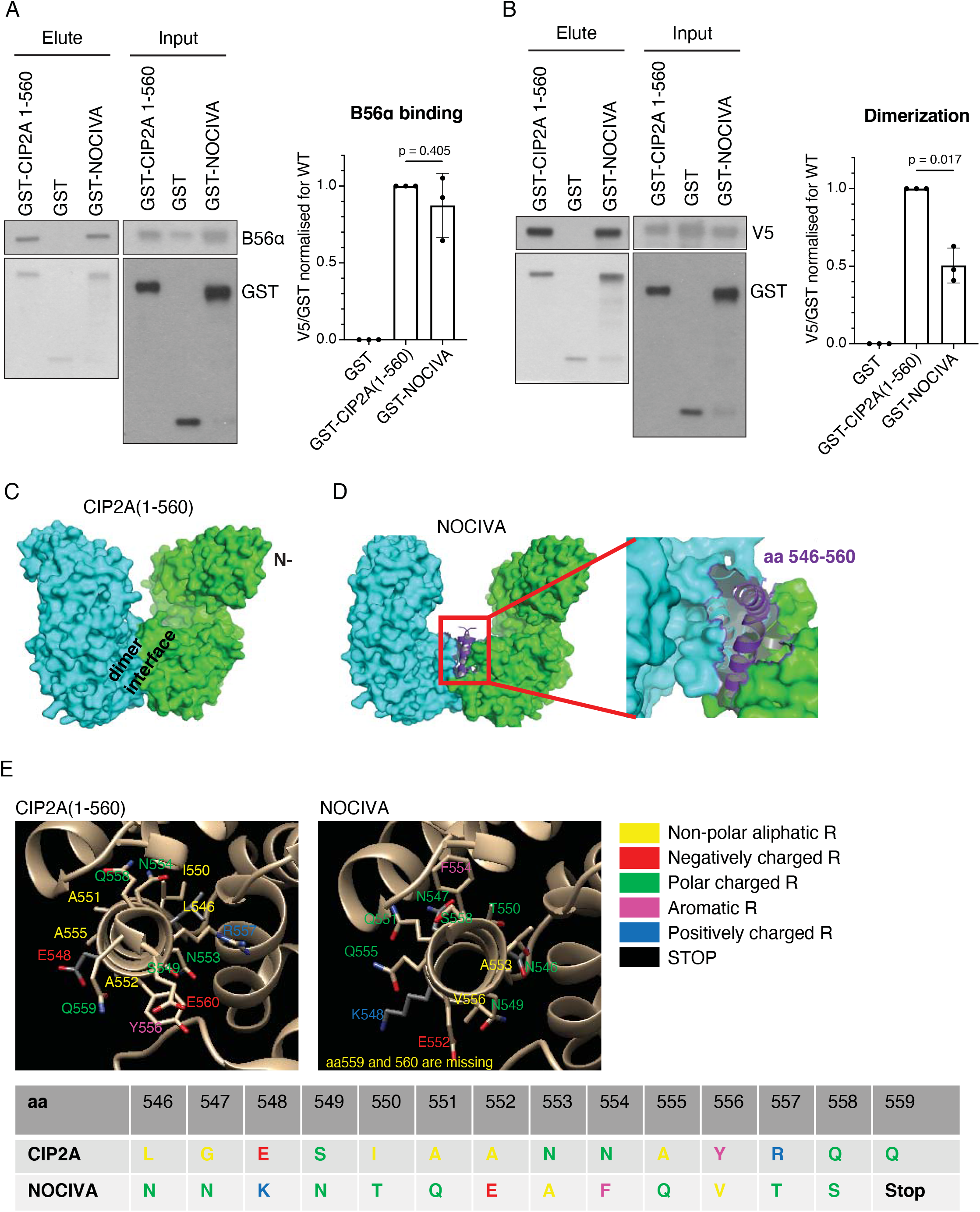
Characterization of NOCIVA protein. **A)** *In vitro* GST-pulldown assay for interaction between PP2A B56α and GST-CIP2A(1–560), and GST-NOCIVA. Equal molar amounts of GST, GST-CIP2A(1–560) and GST-NOCIVA proteins were incubated with B56α for 1 h at 37°C before pulldown. Representative images from three experiments are shown. The accompanying graph shows relative B56α-binding efficiency of GST-NOCIVA as compared to GST-CIP2A(1–560), quantified as a ratio between B56α and GST-CIP2A(1–560) in pulldown samples. Each bar is mean ± SD from three independent B56α-binding experiments; p = 0.405 by one sample t-test. **B)** *In vitro* hetero-dimerization assay using purified recombinant GST-tagged NOCIVA and CIP2A(1–560) proteins. Equal molar amounts of GST, GST-CIP2A(1–560) and GST-NOCIVA proteins were incubated with CIP2A(1–560)-V5 fragment for 1 h at 37°C before pulldown. Samples were analyzed by Western blot using V5 and GST antibodies. Representative images from three experiments are shown. The graph shows relative dimerization efficiency of GST-NOCIVA as compared to GST-CIP2A (1–560), quantified as a ratio between CIP2A(1–560)-V5 and GST-CIP2A(1–560) in pulldown sample. Each bar is mean ± SD from three independent experiments; p = 0.017 by one sample t-test. **C)** CIP2A exists as obligatory dimer. Crystal structure of CIP2A(1-560) (PDB: 5UFL) shown as surface representation, with indicated dimer interface. Individual monomers in cyan and green. **D)** Zoom into dimer interface area modified in NOCIVA (modified part is in red line). Differences in NOCIVA (residues 546-560), in contrast to CIP2A, are mapped on CIP2A’s surface and shown in purple-blue and as transparent surface representation. Same orientation like in E. E-F were generated in Pymol. **E)** Amino acid residues distinct between CIP2A(1-560) (left panel) and NOCIVA (right panel) are indicated as sticks and colored based on heteroatom. Protein molecule orientation was held in approximately the same for both panels but twisted slightly to show the optimal orientation of the key residues. In CIP2A-NOCIVA dimer stabilizing hydrogen bonds and salt bridges between S519-N553-R557-D520-Y556 and Q559-E560 are lost as compared to CIP2A-CIP2A dimer. Image was done using UCSF Chimera1.14. Differences in the nature of amino acid side chains are represented by the color scheme and also indicated in the alignment, following the same coloring pattern.

Together, these results identify NOCIVA as a novel nuclear CIP2A variant protein, that can heterodimerize with CIP2A and bind directly to the B56α tumor suppressor subunit of PP2A.

### NOCIVA expression in normal and cancer cells

To evaluate the expression levels of *NOCIVA* mRNA in biological samples, and to compare them with *CIP2A*, we designed and validated (details in supplementary methods) two quantitative real time PCR (RQ-PCR) assays for both *NOCIVA* (NOCIVA#1 and #2 assays) and CIP2A (CIP2A e13 and e20 assays). If not otherwise indicated, NOCIVA#1 and CIP2A e20 were the mainstay assays when refering to *NOCIVA* or *CIP2A* mRNA detection in this study.

First, *NOCIVA* and *CIP2A* expression was analyzed in a panel of normal human tissue cDNAs (Human MTC panel I & II, Clontech, cat no 636742 & 636743). Notably, *CIP2A* and *NOCIVA* RQ-PCR assays were optimized to yield similar amplification efficiency allowing direct comparison between their respective expression levels. *NOCIVA* showed overall low levels of expression across normal human tissues (Fig 4A), but consistent with its regulation from the same promoter region than *CIP2A*, expression profile across different tissues, including high expression in testis, was comparable to that of *CIP2A* (Fig. 4A and S5B). To identify tissues in which *CIP2A* AS to *NOCIVA* might be relatively more active we calculated the ratio between *NOCIVA* and *CIP2A* expression across different normal tissues. Although the absolute expression of *NOCIVA* was below 7% of *CIP2A* in all tissues, the leukocytes, kidney, and pancreas were the tissues that had the highest *NOCIVA/CIP2A* ratio (Fig 4B).

**Figure 4.**
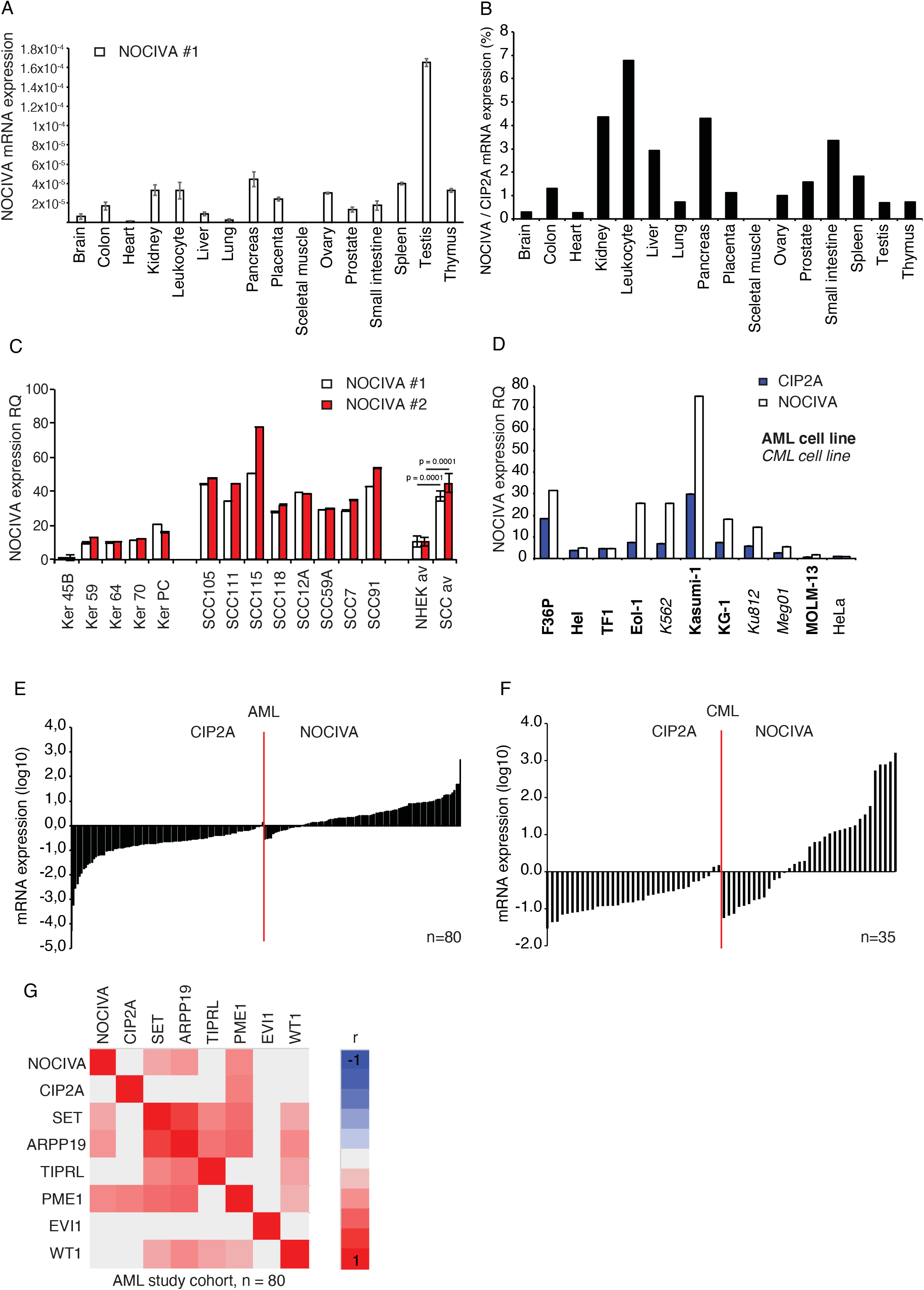
*NOCIVA* mRNA expression in normal and cancerous cells. **A)** *NOCIVA* mRNA expression in normal tissue panel (Human MTC panel I & II, Clontech) measured with NOCIVA #1 RQ-PCR assay. **B)** *NOCIVA* mRNA expression in patient derived normal human epidermal keratinocytes (NHEK, Ker) and squamous cell carcinoma (SCC) cells. mRNA expression levels in Ker 45B cells was defined as value 1. p = 0.0001 by Student’s t-test. **C)** *NOCIVA*/*CIP2A* mRNA expression in normal cells, indicating leucocytes as a tissue for further study. **D)** *NOCIVA* and *CIP2A* mRNA expression in AML, CML and solid cancer cell lines. AML: Acute myeloid leukemia; CML: Chronic myeloid leukemia. mRNA expression levels in Hela cells were defined as value 1. **E)** and **F)** Waterfall plots of analyzed genes from the patient cohorts normalized to the pooled (n=56) normal BM sample. On the y-axis are log10 transformed RQ mRNA expression values derived from two technical replicates in two independent experiments. One bar represents one patient. In the majority of cases, *CIP2A* mRNA expression in AML (E) and CML (F) patient samples was lower than in normal control samples, whereas *NOCIVA* was higher than in normal samples. **G)** Pearson’s pairwise correlations for the mRNA expression of PP2A inhibitors in the AML patient cohort. *NOCIVA* correlates with *PME1* (r=0.43, p=0.0002) and weakly with *ARPP19* (r=0.37, p=0.0014) and *SET* (r=0.30, p=0.0104), but not with other studied markers. Red represents positive and blue negative correlation. Grey indicates non-significant correlation (p-value > 0.05). *Beta-actin* & *GAPDH* were used as housekeeping genes in all experiments presented in this figure. Expression values are derived from three technical replicates in two independent experiments. All the figures show mean ± standard error of mean (SEM).

To address potential overexpression of *NOCIVA* in cancer, we first assessed *NOCIVA* mRNA expression between normal epidermal keratinocytes (NHEK, Ker), and patient-derived head and neck squamous cell carcinoma (HNSCC) cells in which *CIP2A* is overexpressed (Fig S5C)[3, 40]. Interestingly, also *NOCIVA* mRNA showed significantly elevated expression in HNSCC samples as compared to NHEK (Fig 4C, p=0.0001 by Student’s t-test).

Followed by the highest *NOCIVA/CIP2A* ratio in lymphoid cells (Fig. 4B), we next tested whether this preferential *NOCIVA* expression was found also from lymphoid cancer cells. Indeed, relatively higher expression of *NOCIVA* than *CIP2A* was observed in most AML (F36P, Eol-1, Kasumi-1, KG-1, MOLM-13) and CML (K562, Ku812, Meg01) cell lines (Fig 3D). Encouraged by these results, we validated preferential *NOCIVA* gene expression from a panel of clinical 80 AML (BM) and 35 (peripheral blood) CML samples. Consistently with earlier results [21, 31], 96% of AML, and 94% CML, patients, respectively, expressed lower levels of *CIP2A* than normal BM controls pooled from 56 males and females (Fig. 4E,F). However, fully supporting active AS from *CIP2A* to *NOCIVA* in myeloid cancers, 77% of AML, and 65% CML samples displayed overexpression of *NOCIVA* (Fig 4E, F).

AML samples were additionally analyzed for mutual dependencies in gene expression levels between *NOCIVA* and the established AML markers Wilms’ tumor 1 (WT1)[41] and ectopic viral integration site-1 (EVI1)[42]; and the PP2A inhibitor proteins *CIP2A, SET, ARPP19, TIPRL*, and *PME1* [31]. Based on Pearson pairwise correlation analysis, we found that *NOCIVA* expression levels significantly correlated with *PME1* (r=0.43, p=0.0002) and weakly but significantly with *ARPP19* (r=0.37, p=0.0014) and *SET* (r=0.30, p=0.0104), but not with other studied markers (Fig 4G).

These results show that *NOCIVA* has a similar expression pattern across normal human tissues to that of *CIP2A*. However, *NOCIVA* displays robust overexpression in AML and CML in contrast to *CIP2A* underexpression from the same samples.

### Clinical relevance of *NOCIVA* expression in diagnostic AML samples

The results above indicate that the myeloid leukemias AML and CML are malignancies in which active splicing of *CIP2A* to *NOCIVA* is particularly prominent. To understand potential clinical significance of this AS phenomenon, we next analyzed the prognostic significance of *NOCIVA* mRNA expression in 80 AML cases treated with intensive chemotherapy (AML study cohort, [31]). After dividing *NOCIVA* expression into high and low expression according to median (2.18, Q_1_=1.14, Q_3_=6.65), Kaplan-Meier estimates revealed that high *NOCIVA* mRNA expression was a strong indicator of shorter overall survival (OS) (Fig 5A, p=0.022 by log-rank test). Very interestingly, low *CIP2A* (Fig 5B, p=0.073 by log-rank test) expression instead was a borderline significance predictor of longer OS indicating that active AS from *CIP2A* to *NOCIVA* is oncogenic in AML.

**Figure 5.**
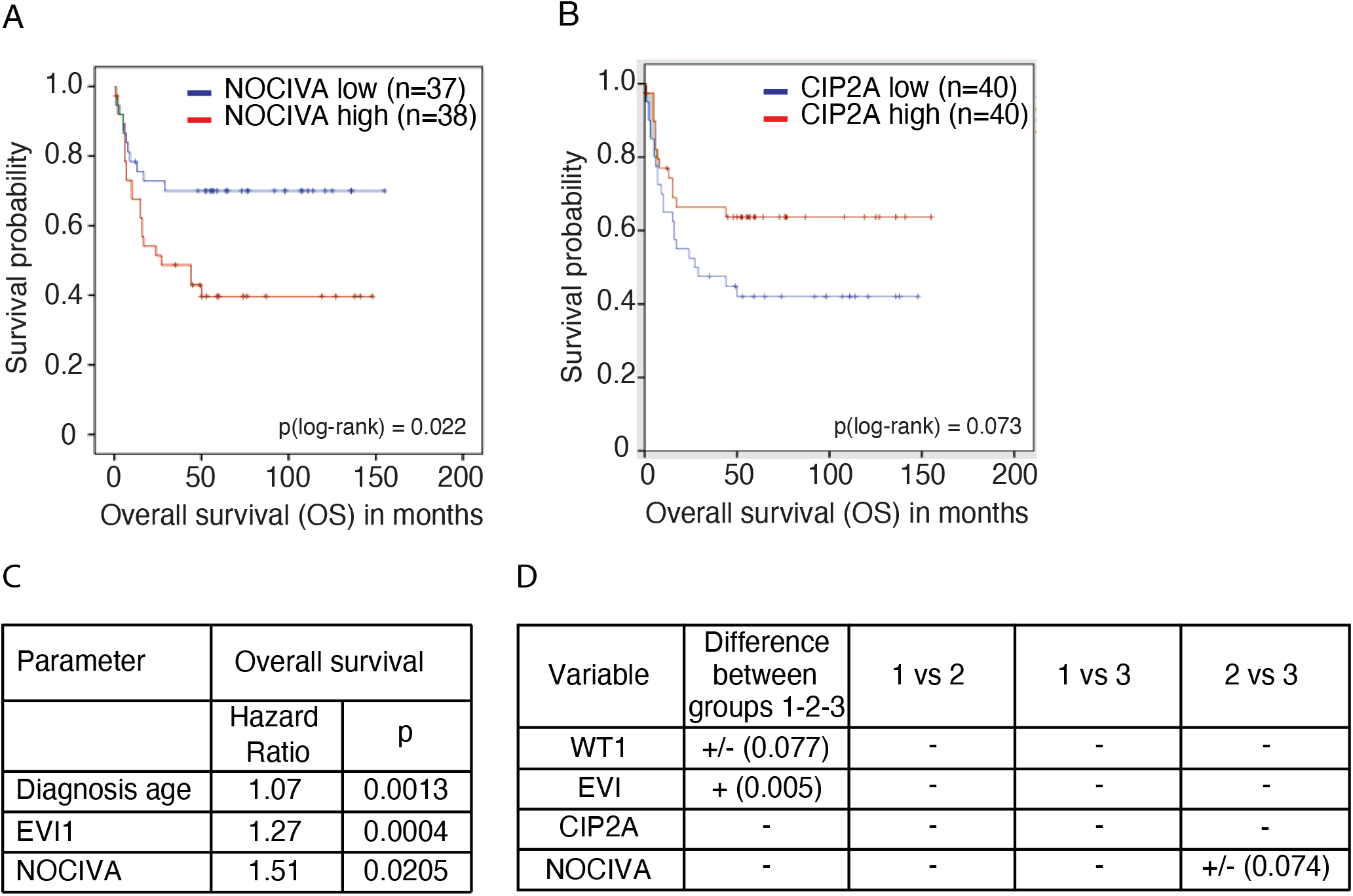
High *NOCIVA* expression is associated with significantly worse overall survival in AML patients. **A)** Kaplan–Meier survival curve for overall survival (OS) by *NOCIVA* gene expression in the AML patient cohort, stratified according to the median expression (see text). Higher *NOCIVA* expression is associated with shorter OS; p = 0.022 by log-rank test. **B)** No significant association was found between *CIP2A* gene expression level and OS (p = 0.073 by log-rank test), though there was a trend towards lower *CIP2A* expression being an indicator of shorter OS. **C)** Multivariable Cox’s proportional hazard model for OS revealed that age at diagnosis (p = 0.0013, HR: 1.07), EVI1 (p = 0.0004, HR: 1.27) and *NOCIVA* (p = 0.0205, HR: 1.51) gene expressions were independent prognostic factors for OS. **D)** Gene expression correlation with risk groups in AML patient cohort by Kruskal-Wallis test. *NOCIVA* expression is a risk group independent prognostic marker in AML. As expected, *EVI1* mRNA expression at diagnosis was significantly different between the three risk groups and its expression increased in relation to the risk group; p = 0.005 by Kruskal-Wallis test. Group 1=favourable, 2=intermediate, 3=adverse. The ELN2010 genetic risk classification was used for risk stratification (see Supplementary Table 2 for detailed information).

Additional analysis for the prognostic role of the studied genes for OS was performed by Cox’s proportional multivariable hazard model, which included age at diagnosis and diagnostic mRNA expression levels of *CIP2A e13, CIP2A e20, SET, ARPP19, TIPRL, PME1, EVI1, WT1* and *NOCIVA*. After excluding the non-significant markers, age at diagnosis (Fig 5C, p=0.0013, hazard ratio (HR): 1.07), EVI1 (p=0.0004, HR: 1.27) and *NOCIVA* gene expression (p=0.0205, HR: 1.51) were found as independent prognostic factors for OS. It was notable that the hazard ratio for *NOCIVA* mRNA expression was even higher than for either *EVI1* expression or diagnosis age which in clinical practise are considered as strong predictors of AML outcome.

We also analysed the association with clinical characteristics and risk groups for the studied markers. The expression of *NOCIVA* or *CIP2A* did not show correlations to any of the clinical characteristics; age, gender, leukocyte or BM blast count, secondary leukemia or the presence/absence of a normal karyotype. In regard to genetic risk group associations, neither *NOCIVA* nor *CIP2A* expression levels showed an association with the ELN2010 risk groups (Fig 5D, p>0.05 by Kruskal-Wallis test). On the other hand, and as expected, *EVI1* mRNA expression at diagnosis was significantly different between the three risk groups, and its expression increased in accordance to the risk group (p=0.005 by Kruskal-Wallis test).

Together these data demonstrate a significant, but risk group independent, association between high *NOCIVA* expression and a poor clinical outcome among AML patients treated with intensive chemotherapy.

### Clinical relevance of *NOCIVA* expression in diagnostic CML samples

Next we evaluated the prognostic significance of *NOCIVA* mRNA expression in 34 newly diagnosed chronic phase (CP) CML patients (CML study cohort1). Twenty patients received imatinib (1G TKI), and 14 dasatinib or nilotinib (2G TKI), as the first line therapy. As calculation of OS was not reasonable in this cohort due to only one death at 60 months, Kaplan-Meier estimates were used to analyze the event free survival (EFS). Importantly, after dividing NOCIVA expression into high and low expression according to median (5.5, Q_1_=0.20, Q_3_=20.0), analysis revealed that high *NOCIVA* mRNA expression was associated with significantly shorter EFS (Fig 6A, p=0.024 by log-rank test).

**Figure 6.**
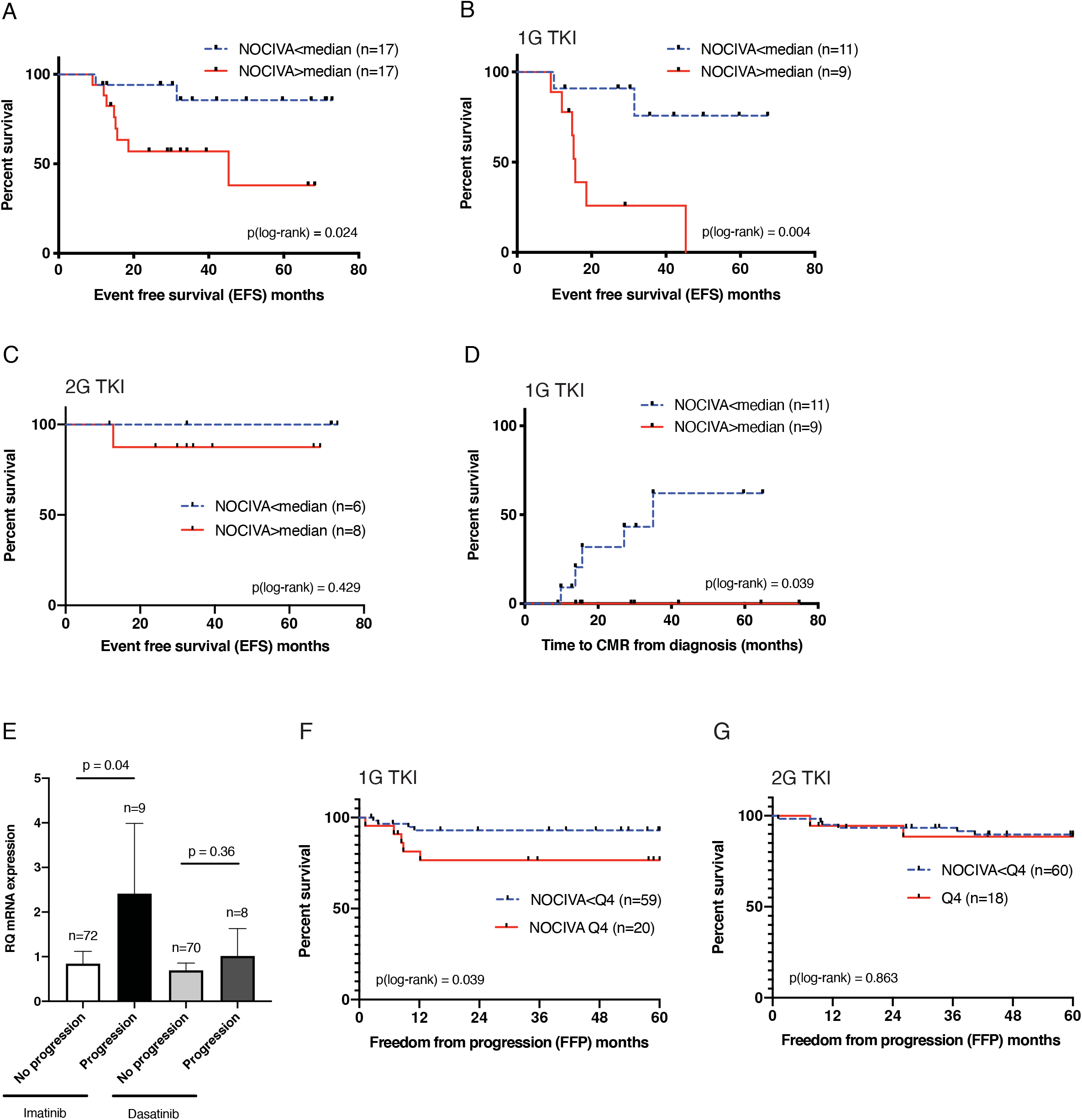
High *NOCIVA* expression is associated with significantly worse outcome, including disease progression, in CML patients treated with imatinib. **A)** Kaplan–Meier survival curve for event free survival (EFS) by *NOCIVA* mRNA expression in CML patient cohort1. Higher *NOCIVA* expression is associated with shorter EFS; p = 0.024 by log-rank test. **B)** Higher *NOCIVA* expression is associated with shorter EFS in imatinib treated patients in CML cohort1; p = 0.004 by log-rank test. **C)** No significant association was found related to *NOCIVA* gene expression level and EFS in patients treated with 2G TKI in CML cohort1; p = 0.429 by log-rank test. **D)** Lower *NOCIVA* mRNA expression is associated with shorter time to complete molecular response (CMR) in imatinib treated patients in CML cohort1; p = 0.039 by log-rank test. **E)** High *NOCIVA* expression at diagnosis is associated with disease progression for imatinib-treated patients in CML patient cohort2; p = 0.04 by Mann-Whitney u-test. Data represents mean ± SEM. **F)** High *NOCIVA* expression is associated with shorter freedom from progression (FFP) in imatinib treated patients in CML cohort2; p = 0.039 by log-rank test. Q_4_ = highest quartile of *NOCIVA* mRNA expression. **G)** No significant association was found related to *NOCIVA* gene expression level and FFP in patients treated with dasatinib in CML cohort2; p = 0.863 by log-rank test. Q_4_ = highest quartile of *NOCIVA* mRNA expression. 1G = first generation, 2G = second generation, TKI = tyrosine-kinase inhibitor. The median level of NOCIVA mRNA expression was used to define the high and low groups in each panel.

Very interestingly, EFS was significantly shorter in the high *NOCIVA* patient group treated with imatinib (Fig 6B, p=0.004 by log-rank test), but this was not seen among the patients treated with 2G TKI (Fig. 6C, p=0.429 by log-rank test). Time to Complete Molecular Response (CMR) analysis was used to assess the depth of a patient’s response, with CMR being the deepest form of response. Patients with high *NOCIVA* expression had a significantly inferior time to CMR (Fig 6D, p=0.039 by log-rank test). Critically, no patient with high levels of *NOCIVA* mRNA at diagnosis achieved CMR. Again, among the patients treated with 2G TKI’s, no association was found between *NOCIVA* expression and CMR, indicating that 2G TKI therapy may overcome the adverse effect of high mRNA expression of NOCIVA.

These findings were then validated in the 159 patient independent CML study cohort2 from the SPIRIT2 clinical trial [33]. In this cohort, 81 patients had received imatinib and 78 dasatinib as the first line therapy. As seen in the CML study cohort1, also here high *NOCIVA* expression at diagnosis was associated with disease progression only among the imatinib-treated patients. Imatinib-treated patients who subsequently progressed to blast crisis had significantly higher expression of *NOCIVA* at diagnosis, than those patients who did not progress (Fig 6E, p=0.04 by Mann Whitney u-test). No significant difference was observed for those patients treated with dasatinib (Fig. 6E). Interestingly, imatinib-treated patients with highest quartile *NOCIVA* expression at diagnosis had significantly inferior freedom from progression (FFP) as compared to patients with lower *NOCIVA* expression (Fig 6F, p=0.039 by log-rank test). Consistent with results from CML study cohort1, no association between *NOCIVA* expression and FFP was observed among the dasatinib-treated patients (Fig 6G).

In conclusion, high NOCIVA mRNA expression assessed at CML CP diagnosis is associated with an inferior EFS and FFP as well as lower rates of CMR selectively among the imatinib-treated patients.

## Discussion

Cancerous inhibitor of PP2A (CIP2A) is an oncoprotein with clinical relevance in a number of human cancers [2]. Furthermore, CIP2A is a very attractive therapeutic target as it is a direct inhibitor of tumor suppressor PP2A-B56a [1], and has low expression in normal tissues [3]. However, regardless of thorough profiling of CIP2A expression across human cancers, mRNA or protein variants of CIP2A remain uncharacterized. In NCBI The Nucleotide and The Protein database, there are four predictions for *CIP2A* splice variant as well as for protein variants of CIP2A. However, none of the predicted variants resemble *NOCIVA*, and neither their functional nor clinical relevance have been studied. This notion emphasizes both the novelty of the presented work as well as indicates the obvious need for experimental validation of also the predicted *CIP2A* isoforms to comprehensively understand CIP2A regulation and function in cancer.

One of the most interesting features of *NOCIVA* is that the intronic region spliced to nucleotide 1636 of *CIP2A*, the last nucleotide of the triplet the codes CIP2A aa 545, is in coding frame with the preceding *CIP2A* sequence and thus codes for a unique, immunogenic C-terminal 13 aa peptide tail (Fig 1B). NOCIVA can be considered as a novel human protein as no sequences homologous could be identified to the 13 aa NOCIVA tail in the human proteome. Strongly indicative of alternative cellular functions of CIP2A and NOCIVA, NOCIVA protein was found to be predominantly nuclear whereas CIP2A mainly resides in the cytoplasm. However, similar to CIP2A, NOCIVA retains the capability to dimerize and to bind to B56a, indicating that it functions similar to CIP2A as a PP2A inhibitor protein. It is clear that further studies on the differential functional roles of NOCIVA and CIP2A are warranted. Unfortunately, during the project we failed in development of siRNA or CRISPR/Cas9 tools selectively suppressing *NOCIVA*, and therefore rather invested thoroughly in demonstrating clinical relevance of the discovery of *NOCIVA*. It is however clear, that thoroughly validated functional models are needed in the future to untangle the cellular role of *NOCIVA*.

AML and CML patient samples displayed clearly higher expression of *NOCIVA* mRNA over *CIP2A* suggesting for particularly active AS of *CIP2A* in myeloid malignancies. This is interesting as AML and CML are the only cancer types where CIP2A seems to be underexpressed at the mRNA level as compared to normal tissue [21, 22]. We thus postulate that upon leukemogenesis a splicing switch that creates *NOCIVA* from *CIP2A* is activated. As splicing is a highly complex event, also at the *NOCIVA* junction site, exonic splicing silencer sequences (ESS) as well as binding sites for hnRNPs and splice factor can be found (Fig 1C). ESSs work by inhibiting the splicing of pre-mRNA strands or promoting exon skipping by primarily recruiting hnRNP binding. A recent paper reported that the most significant Gene Ontology ‘Processes’ and ‘Networks’ changed in AML blasts compared to normal controls were related to transcription, mRNA processing, and stabilization [44]. They observed changes in the expression of 13 hnRNPs affecting mRNA processing, out of which hnRNP A1, A2B1, C are predicted to bind to the *NOCIVA* junction site. Additionally, the expression of hnRNP K [45], SRSF3 (SRp20) [46] and YB-1 [47] have been shown to be altered in AML, but also to contribute to leukemia progression. Interestingly, SRSF3 [37] and YB-1 [38] are additionally shown to specifically promote exon-inclusion during AS. Detailed analysis of the role of these splice factors in AS of *CIP2A* to *NOCIVA* will be required in the future to better understand regulation of NOCIVA in myeloid cancers.

Our data indicate that high *NOCIVA* mRNA expression associates with poor outcome both in AML and CML. In AML, *NOCIVA* expression was independent of the current genetic risk classification, suggesting that the evaluation of *NOCIVA* expression at diagnosis could provide clinically relevant additional predictive value. We also found that high *NOCIVA* mRNA expression assessed at CML CP diagnosis is associated with an inferior EFS and FFP as well as lower rates of CMR for imatinib treated CML patients. Hence, the data suggest that 2G TKI therapy is required to overcome the adverse effects of high *NOCIVA* expression and that together with other diagnostic biomarkers, detection of *NOCIVA* at CML diagnostic phase might help in treatment decisions between imatinib and 2G TKI.

In summary, this work describes discovery of a novel human gene and protein product with the characteristics of a clinically relevant PP2A inhibitor in myeloid malignancies.

## Acknowledgements

Taina Kalevo-Mattila is acknowledged for superior technical assistance. We thank Maria D. Odero (University of Navarra, Pamplona, Spain) for constructive discussions and advice on the material and methodology related to AML cell lines and CIP2A expression in AML. We also gratefully acknowledge the CML subgroup of the United Kingdom National Cancer Research Institute, especially Prof. Jane Apperley, and Sandra Loaiza for access to the SPIRIT2 CML samples and Newcastle University for supplying data from the SPIRIT2 trial. We want to thank Otto Kauko and Juha Okkeri for constructive discussions and advice on metholodology in the beginning of this project. This work was supported by funding from The Sigrid Juselius Foundation, Turku Doctoral Program of Molecular Medicine, University of Turku faculty of medicine, Turku University Hospital ERVA (13283), The Päivikki and Sakari Sohlbergin Foundation, The Cancer Foundation Väre, The Finnish Concordia Fund and Business Finland TUTL (2445/31/2017).

## Authorship Contributions

E.M. and J.W. conceived the study and experiments; E.M., K.P., T.V., S. N., V.K.B. and C.L. performed the experiments; E.M., E.L. and C.L. analyzed the data; U.S. and M.I-R. collected samples and data from AML patients; R.E.C and C.L collected samples and clinical data from CML patients; V-M. K. provided HNSCC & NHEK cDNA panel; E.M. wrote the manuscript, with input from J.W., K.P., U.S., C.L., R.E.C. and M.I-R. All authors reviewed and approved the final manuscript.

## Disclosure of Conflicts of Interest

J.W. and E.M have patents pending for “A NOVEL CIP2A VARIANT AND USES THEREOF” (PCT/FI2018/050844) and “METHOD FOR PREDICTING RESPONSE TO TREATMENT WITH TYROSINE KINASE INHIBITORS AND RELATED METHODS” (PCT/FI2020/050257). In the past 3 years, R.E.C and C.L. declare relevant research support from Bristol Myers Squibb, and R.E.C. declares non-relevant research support and honoraria from Pfizer and Novartis and non-relevant honoraria from Jazz pharmaceuticals and Abbvie.

---

**Supplemental Figure 1.**
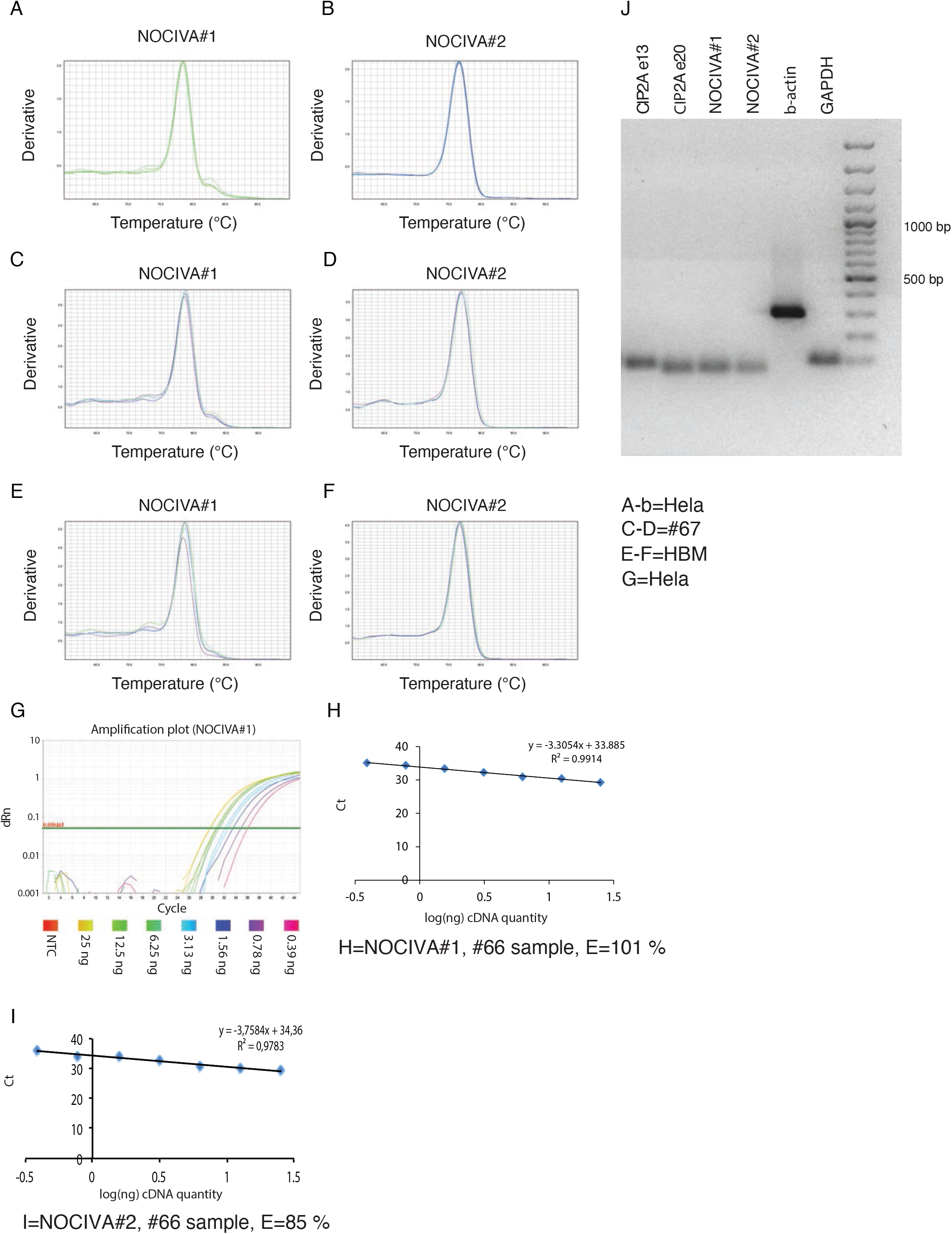

**Supplemental Figure 2.**
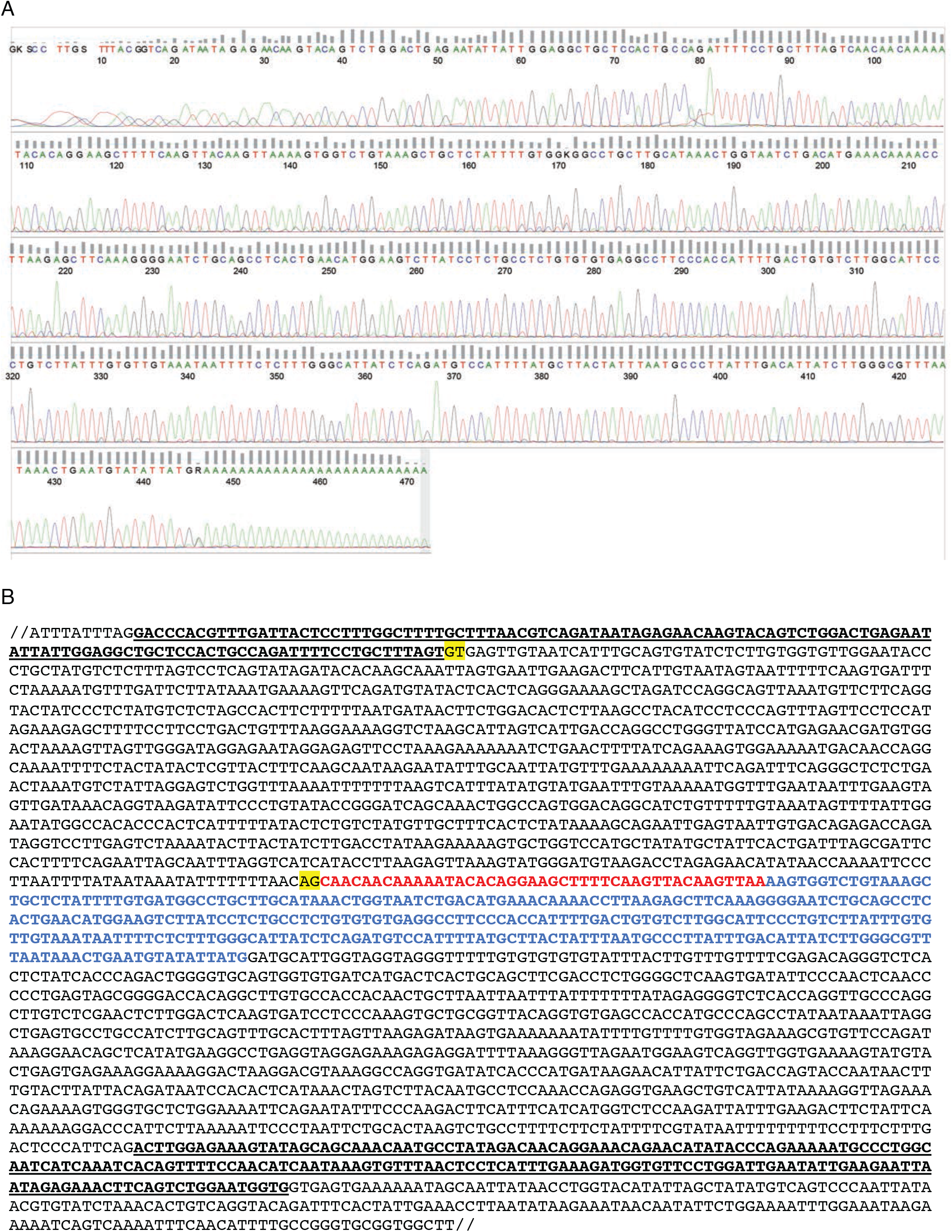

**Supplemental Figure 3.**
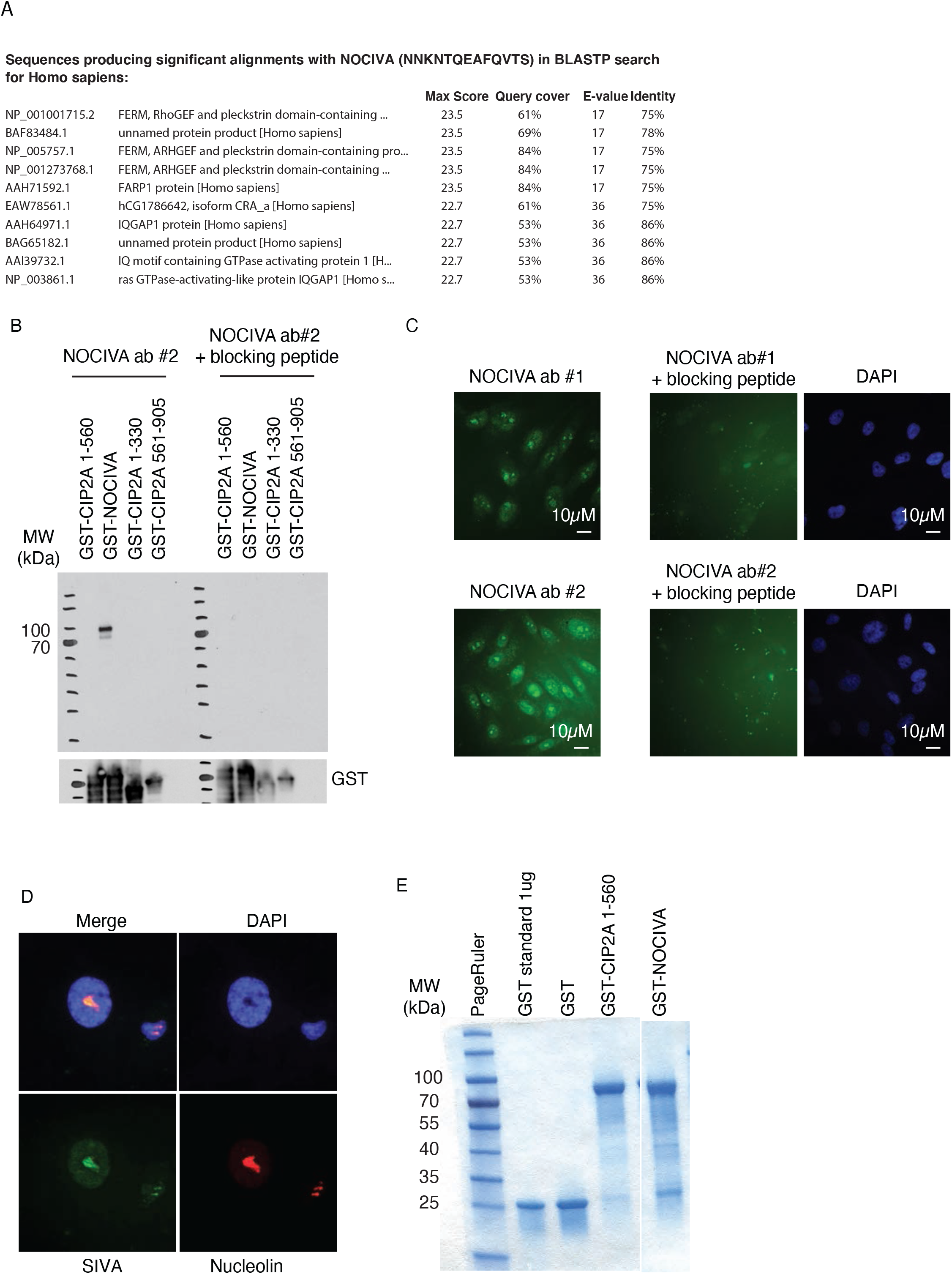

**Supplemental Figure 4.**
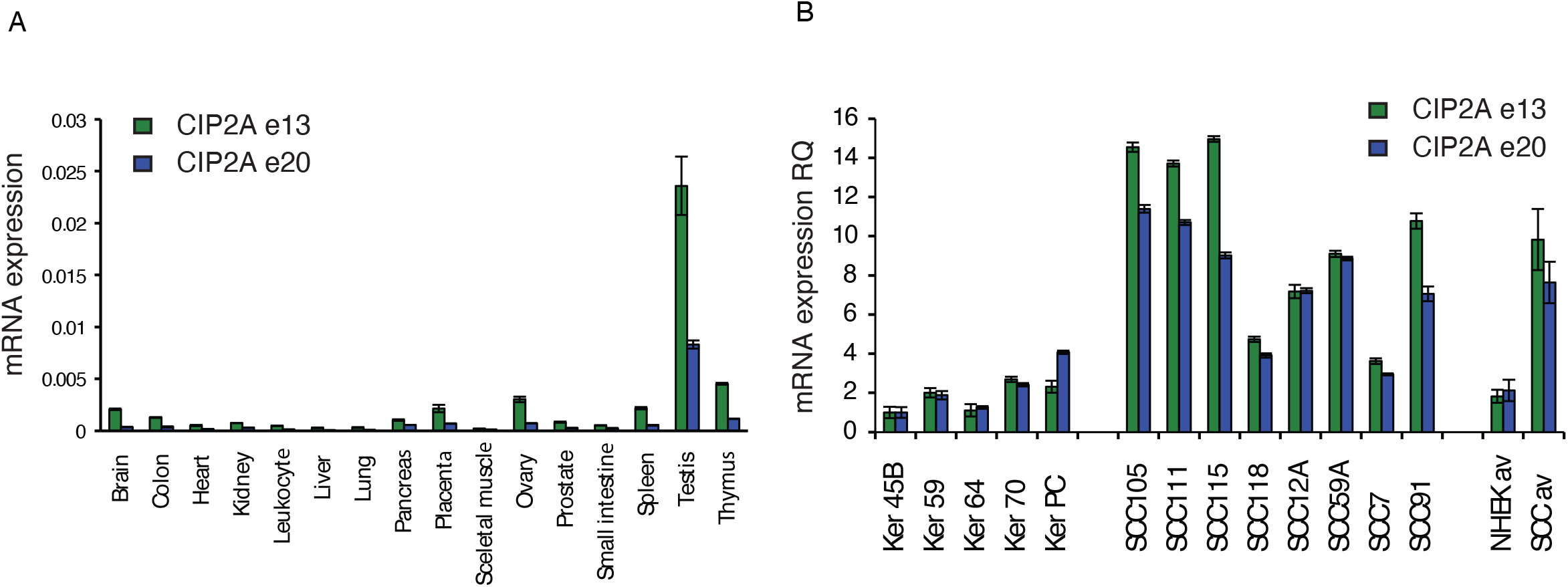

**Table S1.**
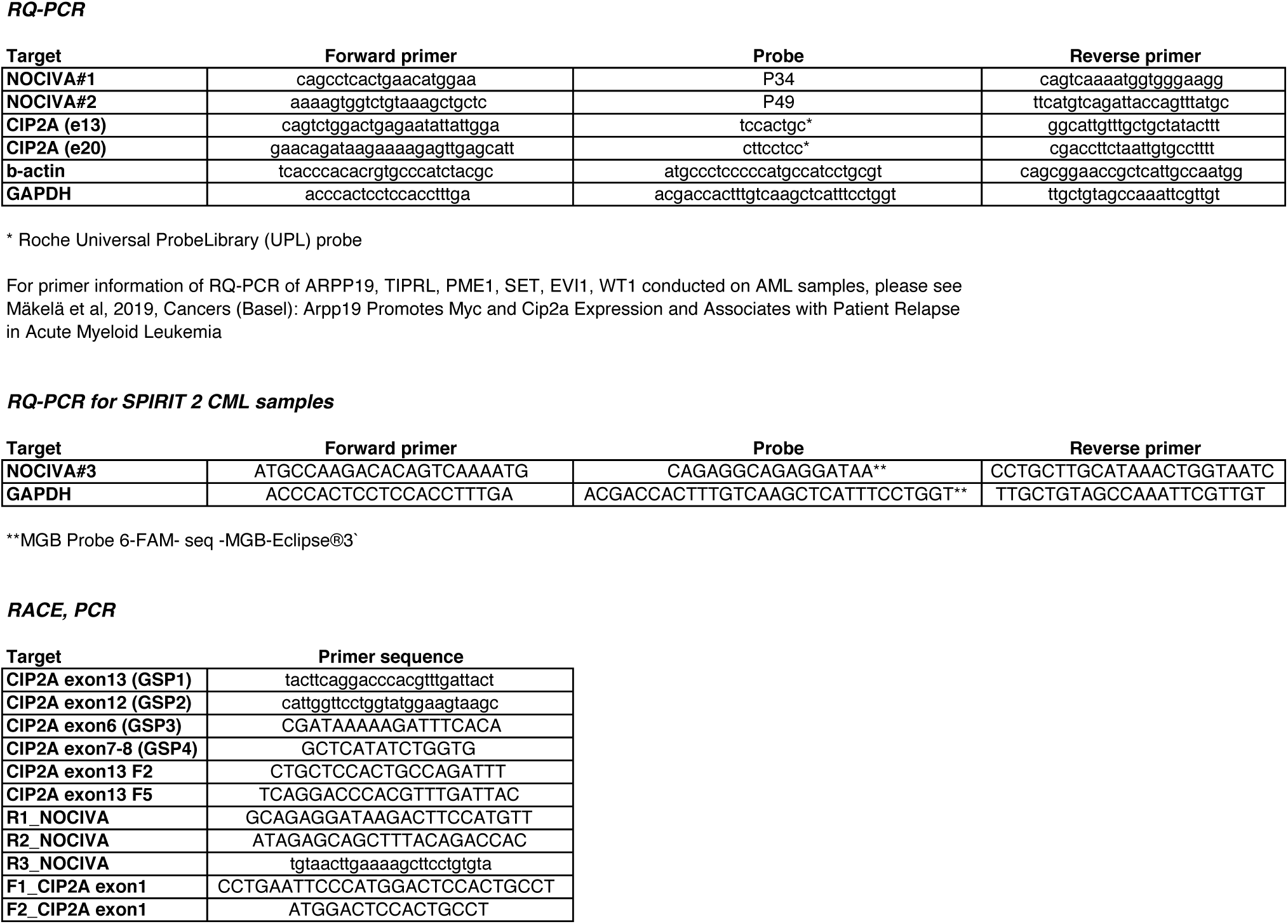
Primer and probe sequences used in this study for PCR based analysis

